# Targeting translation activity at the ribosome interface with UV-active small molecules

**DOI:** 10.1101/436311

**Authors:** Divya T. Kandala, Alessia Del Piano, Luca Minati, Massimiliano Clamer

## Abstract

Puromycin is a well-known antibiotic that is used to study the mechanism of protein synthesis and to monitor translation efficiency due to its incorporation into nascent peptide chains. However, puromycin effects outside the ribosome catalytic core remain unexplored. Here, we developed two puromycin analogues (3PB and 3PC) that can efficiently interact with several proteins involved in translation, ribosome function and RNA processing. We biochemically characterized the binding of these analogues and globally mapped the direct small molecule-protein interactions in living cells using clickable and photoreactive puromycin-like probes in combination with in-depth mass spectrometry. We identified a list of proteins that interact with ribosomes during translation (e.g. eEF1A, ENO1 and GRP78) and we addressed possible uses of the probes to sense the activity of protein synthesis and to capture associated RNA. By coupling genome-wide RNA sequencing methods with these molecules, the characterization of unexplored translational control mechanisms will be feasible.

## INTRODUCTION

The ribosome is a dynamic platform that converts genetic information within mRNA into corresponding polypeptide sequences. Many aspects of this translation machinery have been revealed, from the characterization of ribosome structure and catalytic activity^1,2^ to the organized structure of polyribosomes.^3,4^ At the very beginning of these studies, many antibiotics were used to probe the fundamental mechanisms of protein synthesis. Puromycin, an analogue of the 3’-end of tyrosylated-tRNA,^5,6^ was one of the first drugs used for this application. Puromycin participates in peptide bond formation^7^ through the irreversible reaction of its α-amino group with the peptidyl tRNA, causing release of the nascent peptide and ribosome dissociation along the transcript.^8^ A number of synthetic derivatives have been tested as substrate analogues to probe ribosome catalytic activity, to control puromycin incorporation in nascent peptides^9–12^ and to sense intracellular chemical compounds.^13,14^ Nevertheless, the activity of puromycin derivatives outside the framework of peptide bond formation has never been examined in detail. Here, we show that α-amino-modified and UV-active puromycin derivates, can be used to study protein synthesis activity and label productive ribosome-associated protein.

## EXPERIMENTAL SECTION

### *Syntheses of 3PA, 3PB and 3PC* compounds (collectively called 3xsPx)

A solution of BOC anhydride in CHCl3 was added to a solution of diethanamine to provide a mono-NBOC derivative, which was then reacted with an equimolar amount 3-(prop-2-ynyloxy)propanoic acid. The clean product was deprotected, collected and treated dropwise with an equimolar amount of CDI in pyridine. Puromycin was then added to the CDI-activated molecule at 100°C for 7 hours.

The crude product was evaporated under vacuum at 50°C, affording a brown oil, which was extracted using different mixtures of AcOEt/water and a DCM/water partitioning system. The organic phase was purified by silica gel column chromatography with DCM/MeOH gradient elution to obtain the pure puromycin-alkyne product. A (diaziridin-3-yl)methyl carbonochloridate solution (155 mg, 1 molar equivalent) in 0.2 mL of DCM was added dropwise over 30 min to a pyridine solution (7 mL) of the puromycin-alkyne derivative (720 mg, 1 molar equivalent) stirred at r.t. under N_2_. The mixture was kept at 0-5°C; after 2 further additions (2 molar equivalents) in 2 h, the reaction was quenched by adding 10 mL of distilled water and 10 mL of AcOEt. The obtained organic phase was dried over MgSO_4_, evaporated at 40°C to give a raw reaction products as an oil, and finally purified by silica gel column chromatography (35 g) with EtOAc/MeOH gradient elution, collecting 120 fractions of 25 mL. Using LC-MS and NMR analyses, fractions 13-30 were determined to contain the bis-aziridino derivative (3PC, 301 mg, 95% purity), fractions 57-90 the mono-aziridino derivative (3PB, 52 mg, 96% purity) and fractions 92-113 the other mono-aziridino derivative (3PA, 275 mg, 97% purity).

*3PA 1^H^ NMR* (400 MHz, Chloroform-d) δ 8.24 (s, 1H), 7.94(s, 1H), 7.18 – 7.10 (m, 2H), 6.94 (d, J = 6.8 Hz, 1H), 6.86 – 6.77 (m, 2H), 6.72 (s, 1H), 6.03 (d, J = 7.5 Hz, 1H), 5.86 (d, J = 2.3 Hz, 1H), 5.66 (s, 1H), 5.50 (t, J = 5.6 Hz, 1H), 4.62 (dd, J = 12.9, 6.2 Hz, 2H), 4.52 – 4.40 (m, 2H), 4.29 (dd, J = 12.0, 5.0 Hz, 1H), 4.21 – 4.10 (m, 3H), 4.06 (t, J = 6.6 Hz, 2H), 3.76 (d, J = 7.5 Hz, 5H), 3.61 – 3.53 (m, 6H), 3.53 – 3.31 (m, 4H), 3.31 – 3.20 (m, 1H), 3.01 (h, J = 6.9, 6.5 Hz, 2H), 2.56 – 2.39 (m, 3H), 1.82 (s, 2H), 1.65 (td, J = 6.6, 2.7 Hz, 2H), 1.04 (s, 3H).

*3PB* 1^H^ NMR (400 MHz, Chloroform-d) δ 8.28 (s, 1H), 7.81 (s, 1H), 7.22 – 7.14 (m, 2H), 6.89 – 6.81 (m, 3H), 6.60 (s, 1H), 6.10 (s, 1H), 5.97 (d, J = 7.6 Hz, 1H), 5.89 (d, J = 5.8 Hz, 1H), 5.76 – 5.69 (m, 1H), 5.44 (t, J = 5.5 Hz, 1H), 4.85 (td, J = 6.8, 4.0 Hz, 1H), 4.50 (q, J = 7.4 Hz, 1H), 4.21 – 3.97 (m, 6H), 3.93 (d, J = 12.9 Hz, 1H), 3.78 (s, 3H), 3.77 (t, J = 5.8 Hz, 2H), 3.70 (s, 1H), 3.64 – 3.55 (m, 6H), 3.52 (t, J = 4.9 Hz, 2H), 3.43 (p, J = 5.8 Hz, 2H), 3.40 – 3.31 (m, 2H), 3.12 – 2.96 (m, 2H), 2.50 – 2.42 (m, 3H), 1.68 (td, J = 6.5, 2.8 Hz, 2H), 1.04 (s, 3H).

*3PC* 1^H^ NMR (400 MHz, CDCl3) δ 8.24 (s, purom,1H), 7.94 (s, purom, 1H), 7.14 (d, 8.6 Hz, purom, 2H), 6.94 (d, J = 6.8 Hz, NHCO, 1H), 6.82 (d, 8.7 Hz, purom, 2H), 6.71 (m, NHCO, 1H), 6.03 (d, 7.5 Hz, COCHNHCO, 1H), 5.66 (brs, ribose CH-OCO, 1H), 5.50 (t, 5.8, NHCO,1H), 4.62 (q, 7.1 Hz, ribose CH-NHCO, 1H), 4.60 (m, riboseCH-1H), 4.46 (dt, 6.1, 8.2 Hz, NHCOCHNHCO, 1H), 4.46 (dd, 12.0, 8.0 Hz, -CH2-OCO, 1H), 4.29 (dd, 12.0, 5.0 Hz, -CH2-OCO, 1H), 4.17 (m, -riboseCH2-OCO,2H), 4.15 (d, 2.4 Hz, -OCH2-alkyne, 2H), 4.06 (m, CH-ribose, 1H), 3.78 (t, 6.2 Hz, -CH2-OCO, 2H), 3.76 (s, CH3)2-N, 6H), 3.65 (t, 6.4 Hz, al-kyneCH2-OCH2-, 2H), 3.52-3.30 (series of m, 8H), 3.46 (s, OCH3 purom, 3H), 3.21 (q, 5.8 Hz, OCH2CH2NHCO, 2H), 3.01 (q, 6.0 Hz, -CH2-NHCONH, 2H), 2.45 (dd, 5.9,13.8 Hz, Ar-CH2-purom, 1H), 2.42 (t, 6.4 Hz, -CH2-CONH, 2H), 2.35(dd, 8.3,13.8 Hz, Ar-CH2-purom, 1H), 1.65 (t, 6.2 Hz, -CH2-aziridine, 2H), 1.04 (s, Me-azir, 3H)

See the Supplementary information for a detailed description of the synthesis and the NMR/MS data.

### Cells and reagents

Breast cancer cell line MCF7 (ATCC catalog no. ATCC^®^ HTB-22™) and HEK-293 (ATCC catalog no. ATCC^®^ CRL-1573™) cells were seeded on adherent plates and maintained at 37°C and 5% CO_2_ in Dulbecco’s modified Eagle’s medium (DMEM) with red phenol supplemented with 10% Fetal Bovine Serum (FBS), 2 mM L-glutamine, 100 units/mL penicillin and 100 μg/mL of streptomycin. Cells were grown to 80% confluence before lysis. Cycloheximide was purchased from Sigma. For starvation, cells at 80% confluence were incubated with DMEM-red phenol supplemented with 0.5% FBS, 2 mM L-glutamine, 100 units/mL penicillin and 100 μg/mL of streptomycin. Cells were kept under starvation for 18 hours at 37°C and 5% CO_2_. Arsenite treatment was performed using cells at 80% confluence, with arsenite added at a final concentration of 1 mM for 1 hour.

### Preparation of cell lysates

MCF7 and HEK-293 were seeded at 1.5 × 10^6^ cells per 100-mm dish and grown until they reached 80% confluence. Cells were washed thrice with chilled PBS containing 10 μg/mL cycloheximide and then harvested with urea lysis buffer (200 mM Tris, 4% CHAPS, 1 M NaCl, and 8 M Urea, pH 8) or hypotonic cytoplasmic lysis buffer (10 mM NaCl, 10 mM MgCl_2_, 10 mM Tris–HCl, pH 7.5, 1% Triton X-100, 1% sodium deoxycholate, 5 U/mL DNase I, 200 U/mL RNase inhibitor, and 10 μg/mL cycloheximide). After hypotonic lysis, nuclei and cellular debris were removed by two centrifugations at 18000 g and 4°C for 5 min. For quantification of the total absorbance value of the cell lysate, the absorbance was measured (260 nm) using a Nanodrop ND-1000 UV-VIS Spectrophotometer before use or storage. The lysates were aliquoted and stored at −80°C.

### Separation nucleus and cytoplasm

The nuclear and cytoplasmic fractions were extracted from MCF7 cells after cycloheximide (10 ug/ml, 5 min, 37°C) and 3PB or 3PBis treatment (10 min, 37°C), followed by irradiation under a UV lamp at 365 nm for 5 min (0.75 J/cm^2^). Cells lysates were prepared as above, and and the pellets containing the nuclei were washed 10 times with PBS to remove cytoplasmic contaminants. Nuclei pellets were resuspended in urea lysis buffer (200 mM Tris, 4% CHAPS, 1 M NaCl, and 8 M Urea, pH 8) and sonicated. The samples were quantified via Bradford Protein assays, and equal amounts of protein were resolved on SDS polyacrylamide gels.

### Cell treatment with 3Px probes

#### Reaction in complete media

MCF7 or HEK-293 cells were grown to 80% confluence and treated or not treated with cycloheximide (10 μg/mL, 5 min, 37°C) and then the probe (10 min, 37°C). Cells were then washed with cold PBS (containing 10 μg/mL cycloheximide), placed on ice and irradiated under a UV lamp (BLX-365, 5 × 8 W) at 365 nm for 5 min (0.75 J/cm^2^), followed by lysis with hypotonic cytoplasmic buffer.

#### Reaction in the cell lysate

HEK-293 or MCF7 cells were treated with cycloheximide (10 μg/mL, 5 min). After hypotonic lysis and precipitation of the nuclei and cellular debris by centrifugation at 18000 g and 4°C for 5 min, the supernatant was diluted to A_260_ = 1 a.u/mL with buffer (10 mM NaCl, 10 mM MgCl_2_, 20 μg/mL cycloheximide, and 10 mM Hepes, pH 7 in DEPC water). 150 μL of thehe diluted lysate was incubated with the reactive probe for 1 hour in a 12-well plate and then UV-irradiated (BLX-365) at 365 nm for 5 min (0.75 J/cm^2^).

### Proteomic analysis

HEK-293 cells or cell lysates were used for LC-MS analysis. *In-column digestion*. Purification and digestion of the chemically bound 3PB target proteins were performed using a Click Chemistry Capture Kit (Jena Bioscience, cat. no. CLK-1065) according to the manufacturer’s instructions. Data from four independent samples were collected. *In-gel digestion of gel slices*. Gel slices were digested using a ProGest digestion robot overnight. Data from four independent samples were collected. *On-bead digestion*. Beads were denatured in a urea buffer composed of urea (6 M) and thiourea (2 M) and then reduced in DTT (10 mM) at 37°C and 1200 rpm for 1 hour. Subsequently, samples were alkylated by adding IAA (at a 50 mM final concentration) in the dark for 1 hour. The magnetic beads were washed with TEAB (100 mM) for 4 times and mixed with the urea buffer. Samples were digested with 1 μg of trypsin in 100 μL of 100 mM TEAB overnight at 37°C with agitation at 1400 rpm. The digested samples were then purified using a Bravo Assay MAP. Data from two independent 3PB bead samples and two PEG bead samples were collected. Magnetic beads were purchased from IMMAGINA Biotechnology (cat. no. 016-00-007-2-1) and GE Healthcare (cat. no. 28-9857-38). *LC-MS*. After tryptic digestion, the samples were resuspended in 40 μL of 2% ACN and 0.05% TFA in HPLC H2O and then 15 μL of each sample were separated by reversed-phase nanoflow HPLC (column: EASY-Spray C18, 50 cm x 75 μm, Thermo RSLCnano; gradient: 0-37 min, 4-30% B; 37-40 min, 30-40% B; 40-45 min, 40-99% B; 45-50 min, 99% B; 50-80 min, 4% B; elution buffers: A = 0.1% FA in HPLC H2O; B = 80% ACN and 0.1% FA in HPLC H2O) and directly analyzed by a Q Exactive HF using: (i) the full ion scan mode m/z range of 350-1600 with a resolution of 60,000 (at m/z 200) and a lock mass m/z = 445.12003 (in-gel samples) and (ii) full ion scan mode m/z range 350-1500 with a resolution of 120,000 (at m/z 400) and a lock mass m/z = 445.12003 (on-bead samples). In-gel and in-column MS/MS was performed using HCD with a resolution of 15,000 on the top 15 ions with dynamic exclusion. On-bead MS/MS was performed using CID on the most abundant ions with a cycle time of 3 seconds and dynamic exclusion.

### Immunoblotting

Cell lysates were prepared as described above. Proteins were separated by SDS–polyacrylamide gel electrophoresis and transferred onto PVDF membranes. Samples were heated at 95°C for 10 min in 6x Laemmli loading buffer and run on SDS-PAGE in buffer containing 25 mM Tris and 192 mM glycine (Bio-Rad, catalog no. 4569033) at 120 V for 50 min. Membranes were blocked with 5% milk (Santa Cruz catalog no. SC-2325) in TBS-Tween (0.1% Tween) for 1 hour, incubated with primary antibody o.n. at 4°C and then washed in TBS-Tween (0.1%, TBST) two times for 10 min each. After incubation with secondary antibodies conjugated to horseradish peroxidase, the blots were washed three times in TBS-Tween (5 min each) and processed using an ECL Plus detection kit (GE Healthcare, Amersham ECL Prime catalog no. RPN2232) or the SuperSignal™ West Femto Maximum Sensitivity Substrate (Thermo Scientific) as instructed by their suppliers.

### Polysome profiling

Cytoplasmic lysates from frozen mouse tissues were prepared as described previously (Lunelli et al., 2016)^15^. Cleared supernatants were loaded on a linear 15%–50% sucrose gradient and ultracentrifuged in a SW41Ti rotor (Beckman) for 1 hr and 40 min at 180,000 g at 4°C in a Beckman Optima LE-80K Ultracentrifuge. After ultracentrifugation, gradients were fractionated in 1 mL volume fractions with continuous monitoring absorbance at 254 nm using an ISCO UA-6 UV detector.

### Immunoprecipitation

For immunoprecipitation, cells were harvested with lysis buffer (10 mM NaCl, 10 mM MgCl_2_, 10 mM Tris–HCl, pH 7.5, 1% Triton X-100, 1% sodium deoxycholate, 0.2 U/mL, 5 U/mL DNAse I, 200 U/mL RNase inhibitor, and 10 μg/mL cycloheximide with protease inhibitor) and precleared with protein A/G Magnetic Agarose beads (Thermo Scientific, catalog no. 78609) for 30 min at 4°C. The precleared lysates were incubated with specific antibodies against p-S6 (240/244), puromycin or the IgG controls for 1 hour at 4°C in buffer (10 mM HEPES, pH 7.5, 150 mM KCl, 5 mM MgCl_2_, and 100 μg/ml cycloheximide). Protein A/G magnetic agarose beads (50 μL, Thermo) were added to individual samples and incubated for 1 hour at 4°C. The beads were then washed extensively with wash buffer (10 mM HEPES, pH 7.5, 350 mM KCl, 5 mM MgCl_2_, 0.05 M DTT, 1% NP40, and 100 μg/ml cycloheximide) on ice. After the final wash, the beads were transferred to new vials, boiled with 6X Laemmli loading buffer for 10 min at 95°C and resolved on SDS polyacrylamide gels, followed by immunoblotting as described above.

### Transient transfection with GRP78-siRNA

For the transient transfection treatment, MCF7 cells were seeded at a density of 3 x 10^6^ cells in a 6-well plate and cultured for 24 hours in DMEM complete medium. When 70% confluence was reached, the cells were transfected for 24 hours with 20 nM GRP78-siRNA (Silencer Pre-designed siRNA, Life technology) using Lipofectamine RNAiMAX reagents (Invitrogen) in Opti-MEM medium (1 mL, Invitrogen) according to the manufacturer’s instructions. Six hours post-transfection, the medium was replaced with fresh complete medium. The GenBank accession number of the GRP78 sequence used to design the siRNA was nm_005347.4, and the siRNA sequence was: sense-5′- GGAAGACAAUAGAGCUGUtt -3′.

### Sub-cellular fractionation

The subcellular fractionation protocol was adapted from Francisco-Velilla et al., 2016. Briefly, MCF7 cell lysates were prepared from two 10-cm culture dishes with 300 μL of lysis buffer. Two consecutive centrifugations for 10 min at 14,000 RPM and 4°C were performed to remove the cellular debris, nuclei and mitochondria. The supernatant was collected (S30 fraction) and then centrifuged at 95,000 rpm (in a TLA100.2 rotor) to obtain the S100 supernatant and the R pellet (ribosomes plus associated factors) fractions. The R pellet was resuspended in 200 μL of high-salt resuspension buffer (5 mM Tris-HCl, pH 7.4, 500 mM KCl, 5 mM MgCl_2_, 2 mM DTT, and 290 mM sucrose), loaded into a 40-20% discontinuous sucrose gradient and centrifuged at 4°C and 95,000 rpm for 2 hours using a TLA100.2 rotor. The obtained pellet (pure ribosomes fraction, RSW) was resuspended, and the supernatant, which contained the dissociated factors, was collected (FSW).

### Copper catalyzed “click” reaction in the cell lysates and pull-down assays

Cell lysates (0.35 mL) were incubated with CuSO_4_ (2.00 μmol), picolyl-PEG_4_-biotin (2.00 μmol) or picolyl-azide-sulfo-Cy3 (Jena Bioscience, 2.00 μmol), THPTA (10 μmol) and sodium ascorbate (100 μmol) overnight a 4°C. For the pull-down experiments, the cell lysates were then incubated with 500 μL of MAGAR-cN (IMMAGINA BioTechnology S.r.l.) for 1 hour at RT.

### Synthesis and purification of 3PB-biotin and 3PC-biotin

3PB or 3PC (1.00 μmol), azide-PEG_3_-biotin (2.00 μmol), CuSO_4_ (2.00 μmol), THPTA (10 μmol) and sodium ascorbate (100 μmol) were dissolved in 1:1 water-DMSO (0.3 mL) and mixed overnight at room temperature. The final product was extracted by CHCl3 with the aid of sonication, and the product was obtained by solvent evaporation under vacuum. The reaction was verified by MS and thin layer chromatography (TLC).

### Bead functionalization with 3PB/3PC-biotin

3PB or 3PC was dissolved in 2 M NaCl and 10 mM Tris, pH 7.5 (in DEPC water), at a final concentration of 1 mM and incubated with the same volume of MAGAR-cN beads for 1 hour. The beads were washed five times with PBS to remove unbound 3PB/3PCmolecules.

## RESULTS AND DISCUSSION

### Design and synthesis

In analyzing the effects of puromycin derivates, we were interested to examine whether the synthetized compound can bind ribosomal proteins or other proteins involved in the synthesis of new polypeptides. We inferred that, for mapping alternative puromycin binding sites in living cells, the probe would need four general features: (i) the ability to cross the cell membrane, (ii) a UV-active moiety to covalently bind to targets, (iii) a latent affinity handle for downstream labeling,^16^ and (iv) a conserved puromycin scaffold for the detection of targets with commercially available antibodies. Based on these considerations, we synthetized three different probes (collectively called 3Px), placing a UV-active diazirine residue at the 5′-OH (3PA), at the 2′-OH (3PB) or at both the 2′- and 5′-OH (celled 3PC) of the ribose. We choose the aliphatic methyl diazirine residue for (i) the small size and minimal steric demands; and (ii) the high specificity: the covalent cross-linking will occur if the molecule is proximal to the target; alternatively, the diazirine will be quenched by water or it will undergo intramolecular rearrangements leading to inactive alkenes.^17^ To avoid puromycin incorporation into new peptide chains, we added an alkyne affinity handle at the puromycin α-amino group using a linker unit (Figure 1a; for details on the chemical synthesis, see the Methods and Supplementary Information sections). The alkyne is used to enrich and identify possible targets via copper-catalyzed azide-alkyne cycloaddition (CuAAC^18^ or copper-catalyzed “click” reactions), while the diazirine moiety on the sugar ring permits covalent binding to the targets upon UV light (365 nm) irradiation.

**Figure 1.**
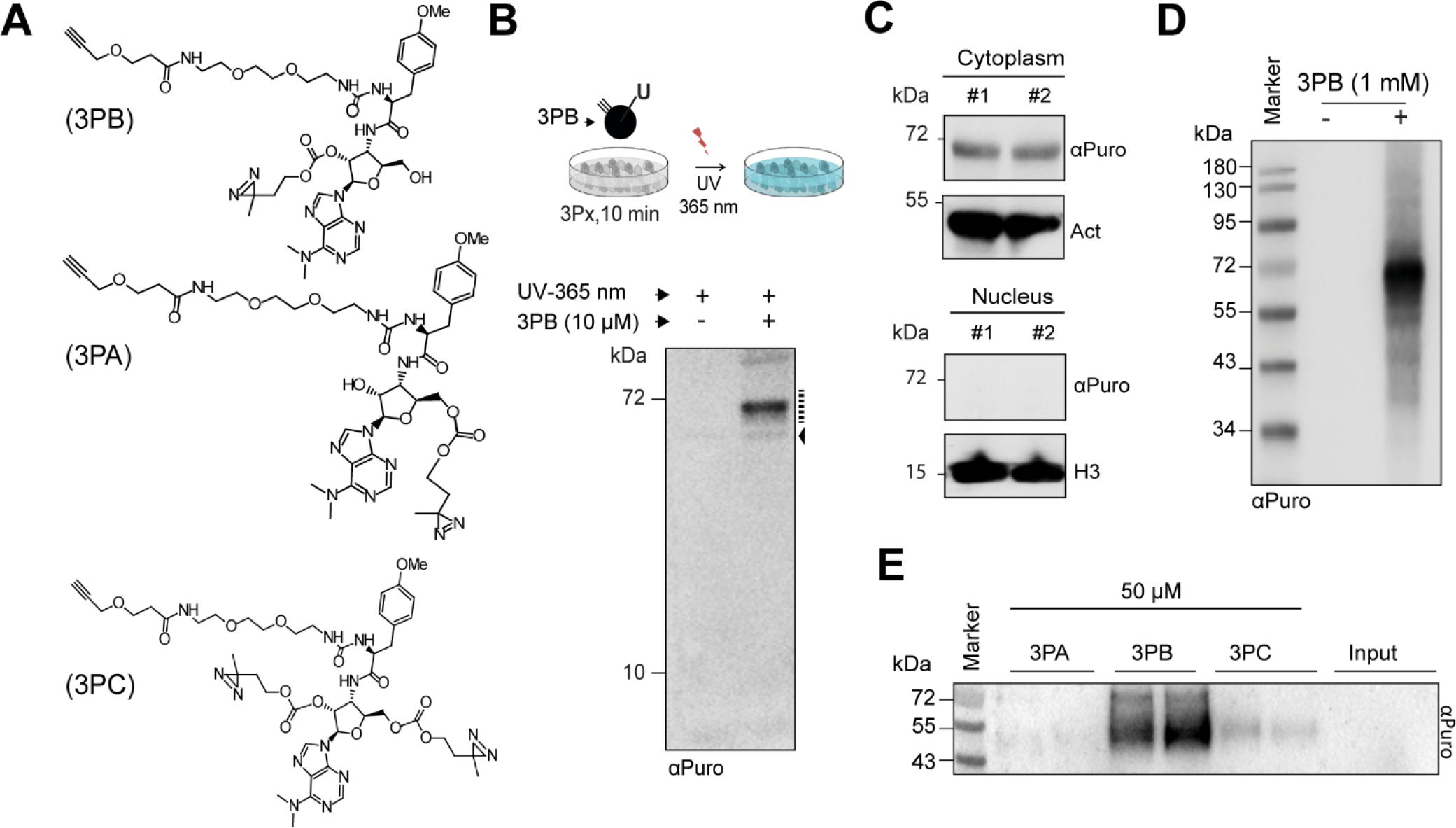
(a) Chemical structure of the 3Px probes. (b, top) Scheme for cell treatment with the 3Px probes. Probes were added to the complete medium and incubated with cells at 37°C for 10 min. After washing with cold PBS, UV irradiation and lysis, the samples were then processed for further analysis. P, photoactive moiety. (b, bottom) Immunoblotting with an anti-puromycin antibody (αPuro) of total protein extracts from urea-lysed HEK-293 cells treated with or without 3PB (10 μM, 10 min). Black broken line, main 3PB targets; black arrow, unspecific signal due to the αPuro antibody. (c) Subcellular fractionation and immunoblotting of MCF7 cells treated with 3PB (100 μM, 10 min). Actin (act) and Histone 3 (H3) were used as cytoplasmic and nuclear markers respectively. #, independent replicates. (d). Immunoblotting with an anti-puromycin antibody of total protein extracts from urealysed HEK-293 cells treated with or without 3PB (1 mM, 10 min). (e) Immunoblotting with an anti-puromycin antibody of cytoplasmic protein extracts from HEK-293 cells incubated with 3PA, 3PB or 3PC (50 μM, 10 min) and exposed to UV (365 nm) irradiation.

### Target profiling in human cells

For our initial analysis, we treated HEK-293 and MCF7 cells in complete culture medium with the 3PB probe (10 μM) for 10 min, followed by UV irradiation for 5 min (total power: 0.75 J/cm^2^). We assessed the labeling of cellular proteins by 3Px using SDS-PAGE analysis. Immunoblotting with an anti-puromycin antibody showed specific puromycin-tagged proteins of approximately 65 kDa in the cytoplasm but not in the nucleus (Figure 1b, 1c and S1a). Increasing the concentration to 1 mM generated multiple bands in the immunoblot, indicating that target specificity decreased at higher concentrations (Figure 1d). The labeling results from UV irradiation, and is concentration- and diazirino–dependent, as demonstrated by comparative experiments with or without UV treatment and with similar molecules lacking the diazirine tag (Figure S1b and S1c). We then compared the labelling activity of 3PB, 3PA and 3PC in HEK293 and MCF7 cells by adding the probes before or after cell lysis. In the latter case, each probe was incubated with cytoplasmic cell extract for 1 hour at 4°C, after which the cell lysates were UV irradiated (0.75 J/cm^2^). We observed a similar pattern to that in the previous experiment, with a concentration-dependent increase in protein labeling for 3PB and 3PC; however, no detectable signals from 3PA were observed (Figure 1e and S1b). Both 3PB and 3PC showed additional bands of ~ 50 kDa when reacted with cell lysate compared to growing cells (Figure 1e and S1d), suggesting a partial loss of selectivity after cell lysis. All these results confirm the binding activities of 3PB and 3PC to few cellular protein targets.

### Protein synthesis activity affects 3Px binding

Because unmodified puromycin binds active ribosomes, we next asked if the binding of our modified puromycin probes was dependent on global translation efficiency. We approached this question by taking advantage of cell treatments known to elicit the repression of protein synthesis (i.e., sodium arsenite and reduced serum concentrations).

We treated MCF7 cells with arsenite (1 mM, 1 hour) or incubated them with 0.5% fetal bovine serum (18 hours). Remarkably, the 3PB protein labelling efficiency was strongly (Figure 2a) reduced by both treatments, suggesting two possible causes: (i) a decrease in the total target protein abundance after stress induction or (ii) different affinity of the probes for the targets upon stress. Therefore, we asked whether the labelling was sensitive to puromycin treatment (i.e., dependent on ribosome activity). To do so, we first treated MCF7 cells with puromycin and then incubated them with 3PB or 3PC for 10 min. When used in minimal amounts, puromycin can be incorporated into neosynthesized proteins,^19,20^ causing ribosome disassembly when incorporated in the growing peptide; conversely, high concentrations of puromycin and long incubation times fully block translation^21^. We defined the concentration that allowed puromycin to be incorporated in the nascent peptide chain (20 μM, Figure S2a). We treated MCF7 cells with both 20 and 100 μM puromycin for 2 hours before adding 3PB (100 μM, 10 min) or 3PC (50 μM, 10 min). Probe-labeled proteins were coupled to a Cy3−azide reporter tag using CuAAC chemistry and were then separated by SDS-PAGE and detected by in-gel fluorescence scanning. We observed a 50% reduction in labelling after puromycin treatment for both probes (Figure 2b and S2b). To better understand if the labelling was related to the activity of the ribosomal peptidyl transferase center (PTC), we pre-treated cells with high concentrations of cycloheximide (CHX, 350 μM) before adding puromycin (100 μM) to fully block ribosome catalytic activity.^9^ Using in-gel fluorescence detection, we observed reduced labelling activity in the CHX-treated sample (Figure 2b and S2b). We confirmed these results by coupling the 3PB-reacted probes to a biotin-azide reporter tag. After SDS-PAGE and membrane staining with streptavidin conjugated to horse-radish peroxidase (HRP), we observed specular effects (Figure S2c). Finally, we investigated if low doses of CHX, which is commonly used to transiently freeze ribosomes along a coding sequence in many gene expression profiles,^22,23^ could affect the labeling. By monitoring the 3PB signal before and after cell treatment with low (10 μM) doses of CHX, we did not observe detectable changes in the intensities of the protein bands (Figure 2c and 2d), suggesting that short (5 min) CHX treatments at a low concentration do not significantly affect labelling. Overall, these results (i) confirm that puromycin alters 3PB/3PC labeling via a mechanism that is independent of the well-known reaction that occurs in the ribosomal catalytic center, (ii) suggest different probe affinities for targets upon translational impairment and (iii) show that CHX) can hamper 3PB/3PC binding only at a high concentration (350 μM)‥

**Figure 2.**
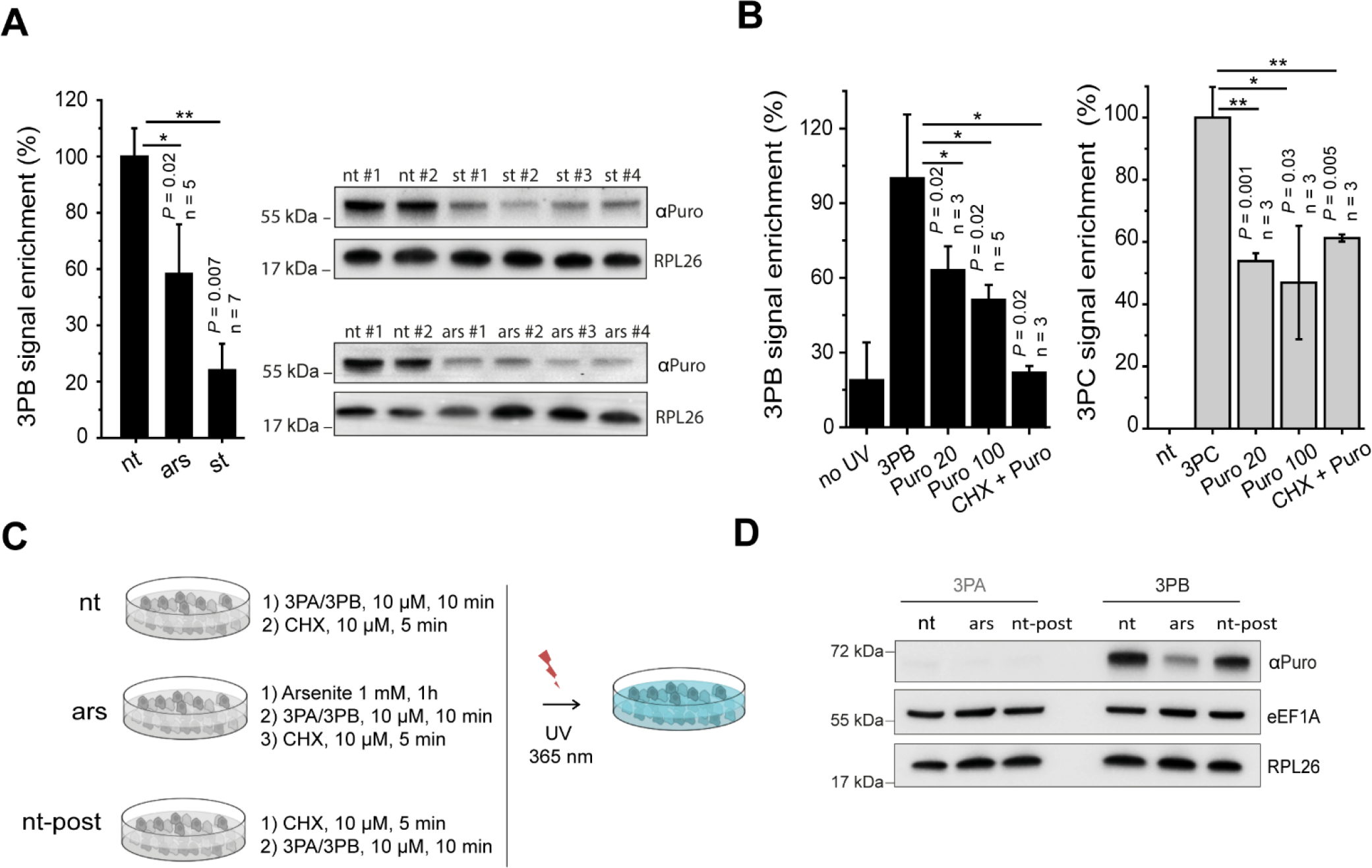
(a) Histogram reporting the pixel quantification of immunoblots of 3PB-tagged proteins at approximately 65 kDa from cells treated with or without arsenite (1 mM, 1 hour) or 0.5% FBS (18 hours). Representative immunoblots are shown on the right. #, independent replicates; nt, not treated; ars, sodium arsenite (AsNaO2) treatment, st, 0.5% FBS. (b) Quantification of the intensity of the 65 kDa bands tagged with 3PB (left) or 3PC (right), with or without UV irradiation and with or without CHX and puromycin added to the cell culture media. Representative images are shown in Figure S2b. Data are normalized (%) to cells treated with 3PB/3PC only. 3PB: 100 μM, 10 min; 3PC: 100 μM, 10 min; Puro 20: 20 μM, 2 hours; Puro 100: 100 μM, 2 hours; CHX+Puro: 100 μM puromycin after treatment with 350 μM cycloheximide. For all experiments, error bars represent the s.d. of three experiments, and t-test P values are reported. (c) Scheme for low-dose cycloheximide cell treatment. MCF7 cells, with (Ars) or without (-) arsenite, were incubated with 3PB (10 μM, 10 min) or 3PA (10 μM, 10 min) before (nt) or after (nt-post) cycloheximide treatment (10 μM, 5 min). After UV irradiation (365 nm, 5 min, 0.75 J/cm2) and lysis with hypotonic lysis buffer, the proteins were quantified and loaded onto an SDS-PAGE gel. (d) Immunoblotting of RPL26, eEF1α and puromycin for 3PA- and 3PB-treated cells according to the scheme reported in (c).

### Identification and analysis of 3Px cellular protein targets

To better understand the responses to inhibitory stresses and drug treatments and to unravel the network of 3PB-target proteins, we used two complementary approaches coupled to in-depth LC-MS studies (Figure S3a). First, we performed an in-column digestion of target proteins after CuAAC chemistry on azide-agarose resin. Second, we pulled down the 3PB-targets with an affinity method based on 3PB-conjugated magnetic-beads, followed by in-gel and on-bead digestion of the captured proteins. Additionally, we used control beads (not functionalized with 3PB) to account for any non-specific signals. In total, 114 protein targets were identified from the in-column digestion (called dataset 1, n = 4), 204 targets from the in-gel digestion (called dataset 2, n = 4) and 1085 from the on-bead digestion (called dataset 3, n = 2, Figure 3a). To increase the stringency of our on-bead digestion dataset, we performed an enrichment analysis using control beads (n = 2) and identified 166 targets (Figure 3a). Six proteins (ENO1, GRP78, eEF1A, PHGDH, KRT2, and HNRNPK) were present in all three MS datasets and at least 2.5-fold more abundant compared to the control beads, suggesting that they are major 3PB targets (Figure 3b and Figure S3b). Interestingly, the detected proteins ENO1, GRP78, eEF1A, PHGDH, and HNRNPK were shown to be associated with ribosomes^24^ and polysomal fractions^25^ in previous mass-spectrometry studies from mouse and human cells. By using the less stringent filter (fold enrichment ≥ 1.3) we identified proteins known to be involved in tRNAs selection and ribosome elongation (eEF2, eEF4G), as well as nascent protein folding (HSP90AB1, PPIA). Only one ribosomal protein (RPL18) was detected in our analysis (fold enrichment of 3PB/Ctrl = 1.4) that is located on the solvent-exposed large ribosomal subunit. These results suggest that the 3PB probe is targeting a complex network of proteins involved with ribosome activity. Among them, we confirm the binding of 3PC for two of the proteins identified in the MS, having ribosome-associated function. The first, a protein involved in RNA binding, ENO1^26^; the second, a folding of the nascent peptide, GRP78.^27–29^ To confirm the binding to the glycolytic enzyme ENO1 and to the chaperone protein GRP78 we performed a selective pull-down assay of the 3PB targets using avidin magnetic beads (Figure 3c). A positive result was obtained for ENO1. For GRP78, we observed the presence of two main protein bands in the input. The first above 72 kDa, and a less intense band between 55 KDa and 72 kDa. GRP78 undergoes post-transcriptional modifications (e.g. ADP-ribosylation and phosphorylation)^29,30^, and two different isoforms of the protein have been described, one that is 72 kDa and a shorter splice variant that is ~62 kDa (GRP78va).^31^ After performing the pull-down assay, we observed the lower GRP78 band, but the upper band (which was dominant in the input) was not detected. In contrast, an immunoblot using an antibody against the parental Hsp-70 protein, which was not enriched in the MS-data, did not show any signal, confirming the specificity of the 3PC pull-down for ENO1 and GRP78. Hence, we silenced GRP78 mRNA expression by introducing GRP78 siRNA into MCF7 cells (Figure S3c and S3d). We found that the 3PC signal after partial (60%) and transient GRP78 knockdown was reduced approximately 50%. Structurally related adenosine-derived molecules are known to inhibit GRP78 as well as its parental protein Hsp70.^32–35^ Since both 3PB and 3PC have adenosine-like moieties, we explored possible competition at the ATP-binding site of GRP78, taking advantage of a specific adenosine-derived inhibitor of Hsp-70, VER-155008. We incubated MCF7 cells with the inhibitor for 1 hour (100 μM) prior to 3PB treatment (100 μM, 10 min). 3PB labelling was strikingly reduced (5.7x) after VER-155008 incubation, suggesting its competitive action with 3PB at the ATP-binding site (Figure S3e). This result confirm the GRP78 isoform as 3PB target. Co-immunoblotting with puromycin and GRP78 on the same membrane showed a common band between 55 and 72 kDa (Figure S3f), which is consistent with the identification of GRP78 as a major 3PC/3PB target. The total protein level of GRP78 did not show evident changes upon serum starvation (i.e., 0.5% FBS, 18 hours), suggesting that 3PB/3PC binds a sub-population of the total GRP78 protein population.

**Figure 3.**
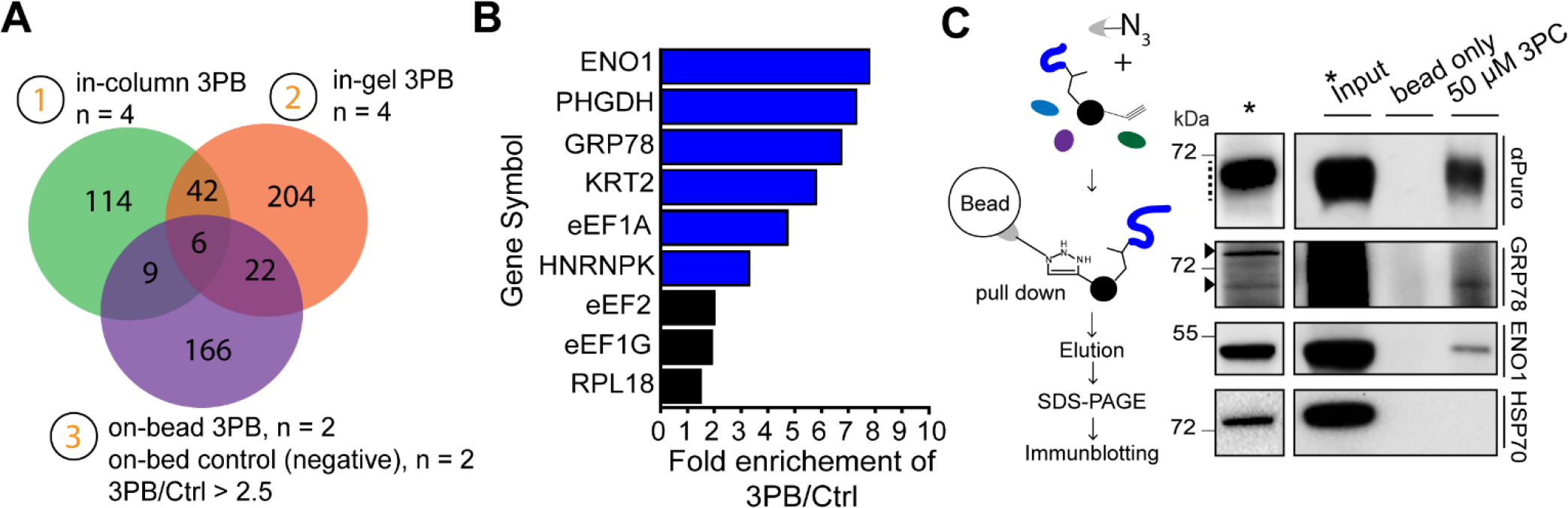
(a) Venn diagram summarizing the proteins detected in groups 1, 2 and 3. (b) Fold enrichment of the top 6 proteins identified (blue bars) and representative proteins less enriched in the LC-MS analysis (black bars). (c, left) Labeling and separation scheme for the 3PB targets via CuAAC reactions for immunoblot detection. (c, right) Immunoblot developed with the indicated antibodies using total protein purified from 3PC-treated cell lysates after CuAAC reactions with a biotin-azide molecule, followed by avidin bead pull-down. Stars, long (right) and short (left) exposure times of the membranes during development.

### Interactions with ribosomes

To confirm the relationships among the putative ~ 65 kDa targets and ribosomes, we used three complementary approaches. First, we performed a sub-cellular fractionation coupled to a high-salt wash to separate the constitutive components of the ribosome from associated factors, according to Francisco-Velilla et al.^36^ The results showed that 3PB-tagged protein associated with the ribosome fraction (Figure 4a), as did elongation factor eEF2, suggesting that the main target is transiently associated with ribosomes. This result was confirmed by the detection of 3PB and 3PC along the polysomal sedimentation profile (Figure 4b and S4). Second, we took advantage of the well-known signature (p240-p244) of the phosphorylated ribosomal S6 protein (p-S6), which is associated with translationally active cells.^37^ We performed a co-immunoprecipitation (co-IP) analysis with an anti-puromycin antibody to investigate the co-precipitation of phosphorylated ribosomal proteins. This analysis revealed the presence of ribosomal protein p-S6 and GRP78 (Figure 4c), confirming that a structural component of the ribosome is an interacting partner. Third, the opposite co-IP with p-S6 showed a strong enrichment in the puromycin signal with respect to the control, IgG, confirming its association with assembled and productive ribosomes (Figure 4d). This result was supported by the evidence of (i) a partial co-localization of RPL26 with Cy3-labelled 3PB/3PC target, as probed by confocal microscopy (Figure S5), and (ii) the sensitivity of 3PC-tagged proteins that co-sediment with the translational machinery to EDTA treatment (which is known to disassemble ribosomes) (Figure S6a). Moreover, the co-IP with the anti-puromycin antibody shows (Figure 4d) the enrichment of PMK2, a glycolytic protein that is known to be an RNA-binding protein for local translation of ER-destined mRNA^24^. An additional co-IP analysis with an anti-Hsc-70 antibody revealed the presence of the puromycin signal together with Hsc-70 (Figure S6b), a chaperone protein known to be associate to the nascent polypeptide^38,39^. All these results suggest the interaction of 3PB/3PC-tag proteins with productive ribosome. Next, we investigated the possibility of using 3PB/3PC to purify associated RNAs. HEK-293 cells were treated with a low concentration (10 μM) of CHX for 5 min to avoid ribosome run-off, and the cell lysates were incubated with 3PB-conjugated magnetic beads (as for the MS-analysis). Total RNA was extracted after being washed and separated, and following quantification, it was found to be 5 times (Figure 4e and S6c) more enriched than the control (non-functionalized beads). Finally, we purified the total RNA from 3PB-beads and we measured by qRT-PCR the relative abundance of the VEGF mRNA, in both high (10% FBS, DMEM) and low (1 mM AsNaO2, 10% FBS, DMEM) performant protein synthesis conditions. We observed a 2-fold enrichment of the transcript on 3PB-functionalized beads in conditions of active translation, with respect to samples treated with arsenite. This variation was specular to the total VEGF protein level (Figure 4f). Overall, these results confirm the binding and separation of total RNA through a protein-mediated ribosome interaction, an essential step for further 3PB/3PC-specific RNA analyses.

**Figure 4.**
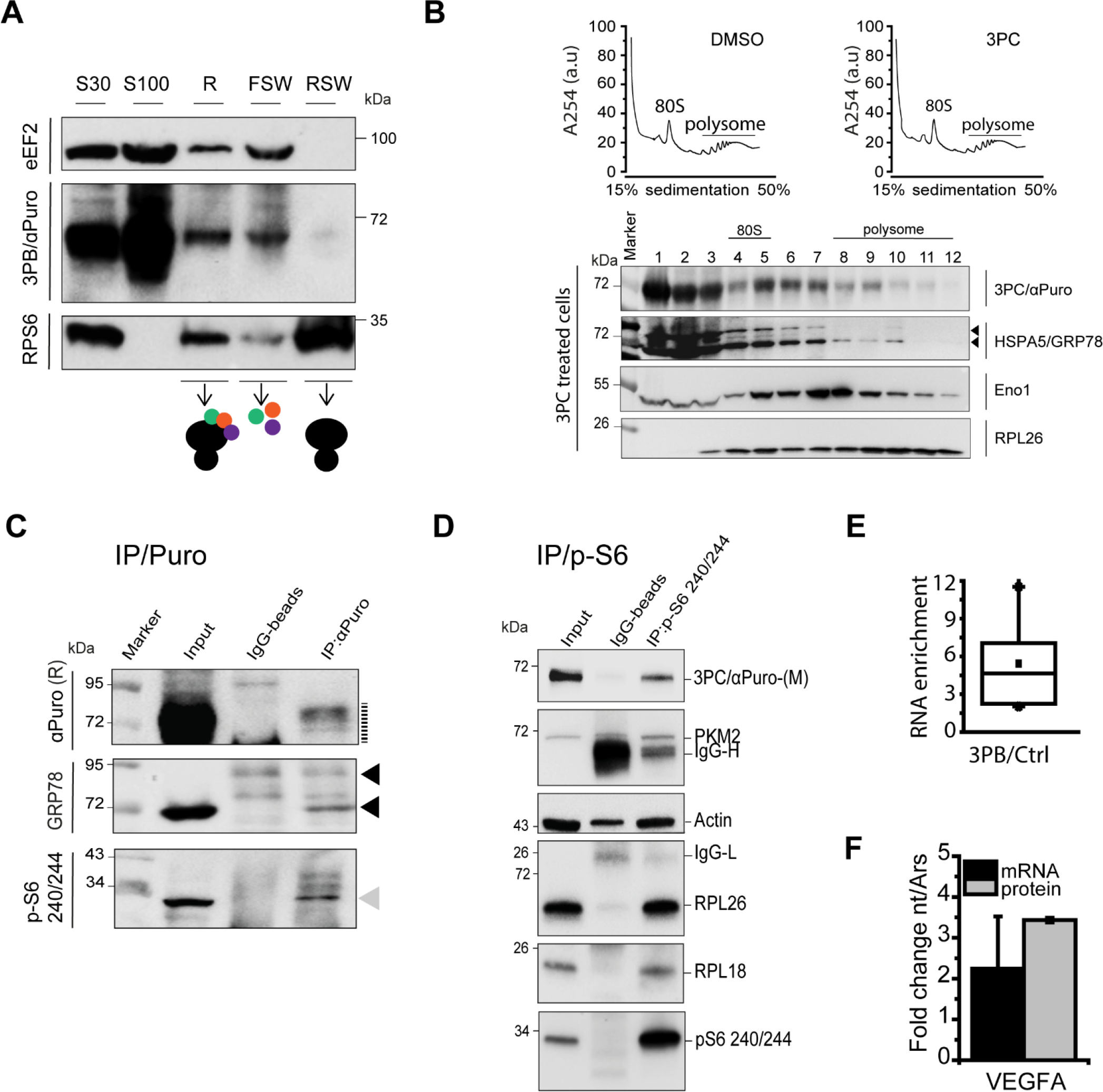
(a) Fractionation and immunoblot analysis of the S30 (input), S100 (soluble), R (ribosomes), FWS (salt-washed ribosomes), and RSW (pure ribosomes) fractions from MCF7 cells after 3PB treatment. The ribosomal protein S6 was used as a control for fractionation. Elongation factor eEF2 was used as a control for RNA-binding protein interactions with ribosomes. (b, top) Sucrose gradient absorbance profile of MCF7 cells at ~ 80% of confluence treated with DMSO or 3PC. (b, bottom) 3PC protein targets co-sediments with the translational machinery. Co-sedimentation profiles of 3PC/αPuro, ENO1, GRP78 and RPL26 by im-munoblotting. Each line corresponds to a sucrose fraction of the profile. Fractions related to the 80S pick or to the polysome region of the profile are indicated. Black arrows, GRP78 main bands. (c) Co-immunoprecipitation analysis of 3PC, GRP78, and p-RPS6. 3PC-tagged protein was immunoprecipitated with a mouse anti-puromycin antibody. Cell lysates before immunoprecipitation (Input) and the immunoprecipitates (IP) were analyzed by SDS-PAGE and immunoblotting (IP) with the indicated antibodies. R, rat anti-puromycin antibody; gray arrow, p-S6. black broken line, 3PB/3PC labeled proteins; black arrows, GRP78 main bands. (d) Co-immunoprecipitation analysis. 3PC-tagged proteins were immunoprecipitated with an anti-p-S6 /240/244 antibody. Cell lysates before immunoprecipitation (Input) and the immunoprecipitates (co-IP) were separated by SDS-PAGE. Immunoblotting was performed with the indicated antibodies. M, mouse anti-puromycin antibody; IgG-H, heavy chain; IgG-L, light chain. (e) Box plot showing the total RNA enrichment on 3PB beads in respect to the control. (f) Histogram with the relative enrichment (no treated and arsenite treated cells) of the VEGFA-mRNA from 3PB-beads (black histogram) and the total VEGFA-protein (grey histogram). Ars, arsenite treatment.

## CONCLUDING REMARKS

Starting from the observation that amino-modified puromycin molecules functionalized with alkyne and diazirine reactive moieties can penetrate cell membranes and bind cytoplasmic protein targets, we demonstrated (i) their selective binding to ribosome-interacting proteins and (ii) their use as sensors of global protein synthesis activity. These findings are supported by three main pieces of evidence. First, we observed that the position of the UV-active (diazirine) moiety on the puromycin molecule is critical for binding. Although all of the molecules synthesized resemble the terminal tRNA, modifications affect their activity: when the UV-active group was placed on the 5′-OH of the ribose (3PA), the molecule was inactive; conversely, when the functional group was on the 2′-OH (3PB) or on both the 5′ and 2′-OH (3PC), the molecules possessed functional activity. Second, a global depression of translation (by puromycin treatment, arsenite treatment or serum starvation) significantly reduced the affinity of the probes for their targets, suggesting that the labeled protein is a distinctive member of the translation machinery. Third, most of the targets identified are known ribosome-interacting proteins.^19,20^ Finally, we demonstrated the binding of 3PB and 3PC to a network of proteins involved in ribosome function and activity (e.g. eEF1A, ENO1, GRP78). In particular, we validated the binding to ENO1 and GRP78. ENO1 is a glycolytic enzyme known to bind ribosomes^24^, eukaryotic tRNA^40^ and specific mRNAs^41^. GRP78 is a protein that (i) controls protein synthesis by regulating eIF2α phosphorylation,^29,42^(ii) participates in nascent protein folding and (iii) contributes to the regulation of cell proliferation, PI3K/AKT signaling and cell viability.^43,44^ Finally, our results showed that a fraction of the targets are not associate to ribosome. Further analysis are needed to accurately address the percentage of 3PB/3PC target associate with ribosomes in vivo. The ability of 3PB and 3PC to sense translation activity (i.e., reduced binding upon depression of protein synthesis), as well as the possibility of efficiently using these probes to purify RNA, paves the way for selective deep-sequencing RNA analysis^45,46^. Overall, these compounds are valuable tools for monitoring protein synthesis and studying translation on the protein-exposed ribosome surface in research, industrial and medical fields.

## ASSOCIATED CONTENT

### Supporting Information

Detailed chemical synthesis of the 3Px molecules with LS-MS/NMR data and schemes. Additional methods and figures reported: pipeline for proteomic MS-data analysis, immunoblotting and antibody descriptions, and Cy3 labeling.

## AUTHOR INFORMATION

### Author Contributions

**MC**conceived the experiments. **DK**and **MC**performed experiments. **ADP**performed the GRP78 silencing experiments. **MC**designed the 3Px synthesis. **LM**assisted with the chemical synthesis of 3Px and performed the synthesis and purification of the 3Px-biotin compounds. **MC**wrote the manuscript. All authors read and approved the final manuscript.

### Notes

MC is a founder, director and share-holder of IMMAGINA BioTechnology S.r.l., a company engaged in the development of new technologies for gene expression analysis at the ribosomal level. ADP, DK and LM are employees of IMMAGINA BioTechnology S.r. IMMAGINA BioTechnology Srl. filed a patent application on 3Px molecules.

## ACKNOWLEDGMENT

We thank **Tocris Bioscience** for assistance with chemical synthesis scale-up, **Manuel Mayr’s Lab** (King’s College London) for providing proteomic LC-MS services, **Gabriella Viero** (Institute of Biophysics, CNR Unit at Trento, Via Sommarive, 18 Povo) for helpful discussion and advice and **Prof. Graziano Guella** (Department of Physics, University of Trento, Povo, Italy) for assistance with the LC-MS and NMR analysis. We thank **Rodolfo F. Gómez-Biagi** for the helpful discussion.

